# Simulation of 69 microbial communities indicates sequencing depth and false positives are major drivers of bias in Prokaryotic metagenome-assembled genome recovery

**DOI:** 10.1101/2023.05.02.539054

**Authors:** Ulisses Nunes da Rocha, Jonas Coelho Kasmanas, Rodolfo Toscan, Danilo S. Sanches, Stefania Magnusdottir, Joao Pedro Saraiva

**Affiliations:** Department of Environmental Microbiology, Helmholtz Center for Environmental Research-UFZ, 04318 Leipzig, Germany; Department of Computer Science, Federal University of Technology -Paraná, UTFPR, Cornélio Procópio 86300-000, Brazil

**Keywords:** Metagenome-Assemble Genome, sequencing depth, metagenomics bias, taxonomic distribution, in-silico microbial communities

## Abstract

We hypothesize that sample evenness, sequencing depth and taxonomic relatedness influence the recovery of metagenome-assembled genomes (MAGs). To test this hypothesis, we assessed MAG recovery in three in silico microbial communities composed of 42 species with the same richness but different sample evenness, sequencing depth and taxonomic distribution profiles using three different pipelines for MAG recovery.

The pipeline developed by Parks and colleagues (8K) generated the highest number of MAGs and the lowest number of true positives per community profile. The pipeline by Karst and colleagues (DT) showed the most accurate results (∼ 92%), outperforming the 8K and Multi-Metagenome pipeline (MM) developed by Albertsen and collaborators. Sequencing depth influenced the accurate recovery of genomes when using the 8K and MM, even with contrasting patterns: the MM pipeline recovered more MAGs found in the original communities when employing sequencing depths up to 60 million reads, whilst the 8K recovered more true positives in communities sequenced above 60 million reads. DT showed the best species recovery from the same genus, even though close-related species have a low recovery rate in all pipelines.

Our results highlight that more bins do not translate to the actual community composition and that sequencing depth plays a role in MAG recovery and increased community resolution. Even low MAG recovery error rates can significantly impact biological inferences. Our data indicates the scientific community should their findings from MAG recovery, especially when asserting novel species or metabolic traits.

## INTRODUCTION

Microbial communities contribute to ecosystems by performing a wide array of ecosystem processes such as the decomposition of organic matter (Tláskal et al. 2021), degradation of contaminants (Huang et al. 2019) or carbon and nitrogen cycling (López-Mondéjar et al. 2018; Soong et al. 2020). The use of metagenomics is a standard method to study microbial ecology, human health and environmental issues, with several databases emerging to facilitate data selection depending on the research questions (Corrêa et al. 2020; Kasmanas et al. 2021; Nata’ala et al. 2022). This use is shown by the exponential increase in deposited Metagenome-Assembled Genomes (MAGs) in public repositories. For example, the National Center for Biotechnology Informtaion (NCBI) started with 132 in 1993 but has more than 1620513 genomes deposited at this time (NCBI, 2023). The Joint Genome Institute (JGI) started with 35 complete or draft genomes in 1997 but currently has 411829 (JGI GOLD, 2023). Advances in Next Generation Sequencing (NGS) and the development of bioinformatics pipelines allow the recovery of near-complete genomes from complex samples, which can be used to characterize microbial communities and assert their functional potential. For example, Parks and colleagues (Parks et al. 2017) recovered nearly 8000 metagenome-assembled genomes (MAGs) using MetaBAT (Kang et al. 2015). Genome annotation of such a large number of MAGs allows to infer the functional potential of species across a diverse array of environmental conditions and provides a wide range of applications. For example, over 12000 studies performed in 2022 rely on MAGs (Dimensions, 2023). The assertions of microbial community composition and their functional profile are thus tightly linked to the reliability of the tools and methods employed to generate MAGs (Haryono et al. 2022). Systematic errors during MAG recovery may profoundly impact the assertions made in microbial community studies that use metagenomics (Meziti et al. 2021). Previous studies have evaluated the effect of individual parameters on genome recovery. Gweon and colleagues (Gweon et al. 2019) showed that sequencing depth affected the profiling of microbial communities while Anyansi and colleagues (Anyansi et al. 2020) highlighted the challenges in the recovery of closely related species. However, studies that simultaneously tackle the different levels of complexity in microbial communities, such as sample evenness and species diversity, are yet to be performed. Minimizing the potential systematic errors during MAG recovery will benefit their growing cascade of ecological and biotechnological applications. With the massive increase in data generated from MAG analyses, several questions remain unanswered; among them: (i) how reliable are the genomes recovered from metagenomes?; (ii) how much of the microbial communities are we missing?; (iii) do the presence of Eukaryote genomes creates bias in the recovery of prokaryotic MAGs?; and (iv) how do sequencing depth, evenness distribution, and taxonomic relatedness between species influence genome recovery?

Reconstruction of MAGs from all domains of life (prokaryotic, eukaryotic and viruses) are mostly performed separately. Pipelines such as those employed by Parks and colleagues (Parks et al. 2017), Sieber and co-workers (Sieber et al. 2018) and Albertsen and colleagues (Albertsen et al. 2013) are just a few examples of prokaryotic MAG recovery. Recent pipelines have been developed for the recovery of eukaryotes (EukRep (West et al. 2018)) and viruses (VirSorter (Roux et al. 2015) and VIBRANT (Kieft et al. 2020)). To our knowledge, only one framework exists that recovers genomes from metagenomes from all domains of life (da Rocha et al. 2022). Independently of the domain of life, much more work needs to be done to increase confidence in the characterization of microbial communities and determine how much of the communities are left uncharacterized. Community diversity and sequencing depth have been suggested to play essential roles in MAG recovery (Meziti et al. 2021). Quality assessment of Prokaryotic MAGs usually relies on sets of single-copy genes (SCGs) or reference genomes to generate results (Milanese et al. 2019). Using only one single-copy gene set may limit the pipelines’ ability to capture all species in a microbial community. For example, strain heterogeneity (Ramos-Barbero et al. 2019) and uneven population abundances of organisms within a community have influenced MAG recovery. The Critical Assessment of Metagenome Interpretation (CAMI) (Meyer et al. 2022) challenges attempts to create benchmarks for comparing metagenomic tools since they are usually difficult to compare. Simulated communities described by Meyer and co-workers (Meyer et al. 2022) mimic some properties of microbial communities, such as multiple closely related strains or species and abundance profiles. Nevertheless, to our knowledge, no dataset exists that contemplates a combination of taxonomic relatedness, sample evenness and sequencing depth and the use of species with multiple or linear chromosomes.

In this study, we tackle MAG recovery knowledge gaps by generating mock communities with varying parameters of sequencing depth, taxonomic distribution and species abundance profiles. We use these datasets to evaluate three pipelines for binning, each using a different approach to bin the sequences, and assess which factors or combinations of factors drive Prokaryotic MAG recovery and if these are pipeline dependent. We also evaluate the effect of multiple or linear chromosomes in MAG recovery.

## RESULTS

### Genome metrics

#### Number of MAGs and quality assessment

The pipelines recovered a total of 2915 (8K), 1429 (DT) and 873 (MM) MAGs from the three communities (Supplemental_Table_S01.xlsx). On average, pipeline 8K recovered the highest number of MAGs in each community profile (42.24, ±122.35), with MM recovering the lowest (12.65, ±27.07). As expected, no eukaryotic genomes were recovered. Average MAG completeness was 96.74 ±4.99 (8K), 97.56 ±4.45 (DT) and 97.32 ±3.49 (MM), and average contamination was 0.363 ±0.505 (8K), 1.25 ±1.65 (DT) and 1.10 ±1.61 (MM) (Supplemental_Table_S01.xlsx and Supplemental_Fig_S1.pdf). On average, the DT and MM pipelines only recovered a single MAG per species, whilst the 8K recovered 3.59 (±0.29) (Supplemental_Table_S02.xlsx).

#### 16S rRNA recovery

The pipelines produced a total of 10 (8K) and 1393 (DT) rRNA gene sequences in the MAGs. Only five 16S rRNA sequences were classified beyond the domain level in MAGs produced by the 8K pipeline. The MAGs produced by the DT pipeline yielded 900 rRNA gene sequences from bacteria, 73 from archaea and 32 from eukaryotes. Only 117, 55 and 16 sequences were classified to family level from each of these domains, respectively (Supplemental_Table_S03.xlsx).

Sankey plots with the species found using the 8K, DT and MM pipelines in all communities (A, B and C) (Supplemental_Table_S04.xlsx) are shown in (Supplemental_Fig_S2.eps – Supplemental_Fig_S10.eps).

### 2.2 Exponential decay has a positive effect on MAG recovery

The 8K pipeline did not yield any significant differences in the number of MAGs recovered between communities in any of the community profiles (evenness distribution *versus* taxonomic distribution (Supplemental_Table_S05.xlsx and Supplemental_Table_S06.xlsx) and evenness distribution *versus* sequencing depth (Supplemental_Table_S07.xlsx).

Using the MM pipeline, communities sequenced at depths greater than 60 million reads yielded significantly lower numbers of MAGs than those sequenced below, independently of the evenness distribution (logarithmic or exponential decay) (Supplemental_Table_S08.xlsx and Supplemental_Fig_ S11.pdf). In “very closely related” communities, a significantly higher number of MAGs (*t-*test, *p* < 0.05) were observed between exponential decay and logarithmic decay evenness profiles (Supplemental_Table_S09.xlsx and Supplemental_Fig_S12.pdf).

When employing the DT pipeline, a significantly lower number of MAGs were obtained in communities sequenced at 10 million reads compared to communities sequenced at 60, 120 and 180 million reads irrespective of evenness distribution (logarithmic or exponential) (Supplemental_Table_S10.xlsx, Supplemental_Fig_S13.pdf).

In “closely related” communities, significantly more MAGs were obtained in communities with an exponential decay evenness distribution compared to communities with logarithmic decay (Supplemental_Table_S11.xlsx, Supplemental_Fig_S14.pdf).

#### 2.2.1 The 8K pipeline recovers the highest number of MAGs

Except for communities sequenced at 30 million reads, the 8K pipeline yielded a significantly higher number of MAGs than the DT and MM pipelines (*t-*test, *p* < 0.05), irrespective of evenness distribution and taxonomic-relatedness (Supplemental_Table_S12.xlsx – Supplemental_Table_S14.xlsx and Figures 1-2).

The DT pipeline yielded a significantly higher number (*t-*test, *p* < 0.05) of MAGs compared to the MM pipeline in communities following a logarithmic decay distribution sequenced at 10, 60 and 180 million reads and in communities following an exponential decay distribution sequenced at 30, 120 and 180 million reads (Supplemental_Table_S13.xlsx and Figure 1).

**Figure 1.**
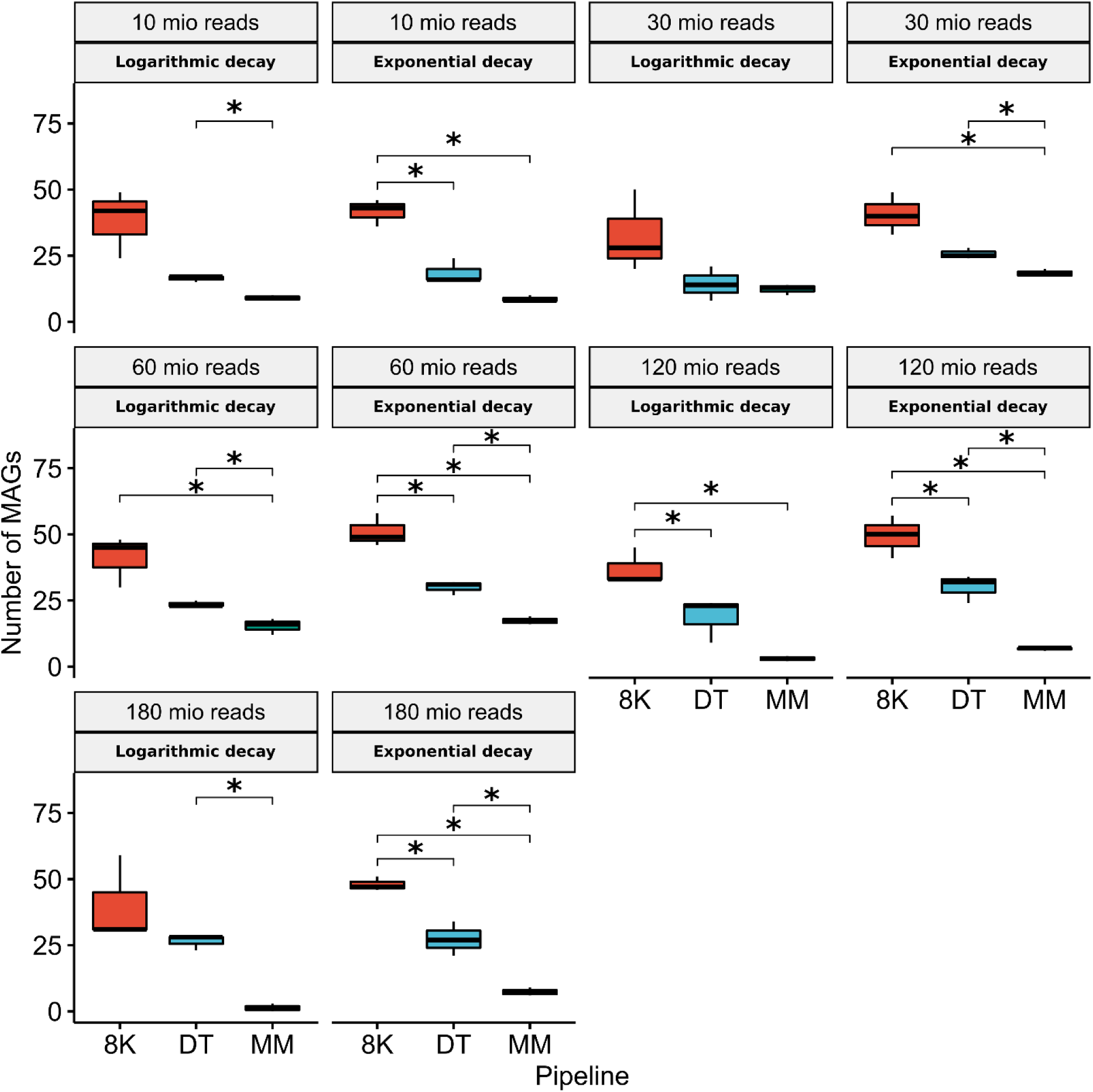
Student’s T-test comparing metagenome-assemble genome counts between all pipelines used (8K, DT and MM) according to evenness distribution (Logarithmic decay; Exponential decay; Logarithmic decay with abundance plateaus; Exponential decay with abundance plateaus) and sequencing depth (10, 30, 60, 120 and 180 million reads). Taxonomic distribution was set to random. (*P-value < 0.05).

Also, the DT pipeline recovered a significantly higher number of MAGs (*t-*test, *p* < 0.05) compared to the MM pipeline in “not closely related” communities following a logarithmic decay with abundance plateaus (Supplemental_Table_S14.xlsx and Figure 2).

**Figure 2.**
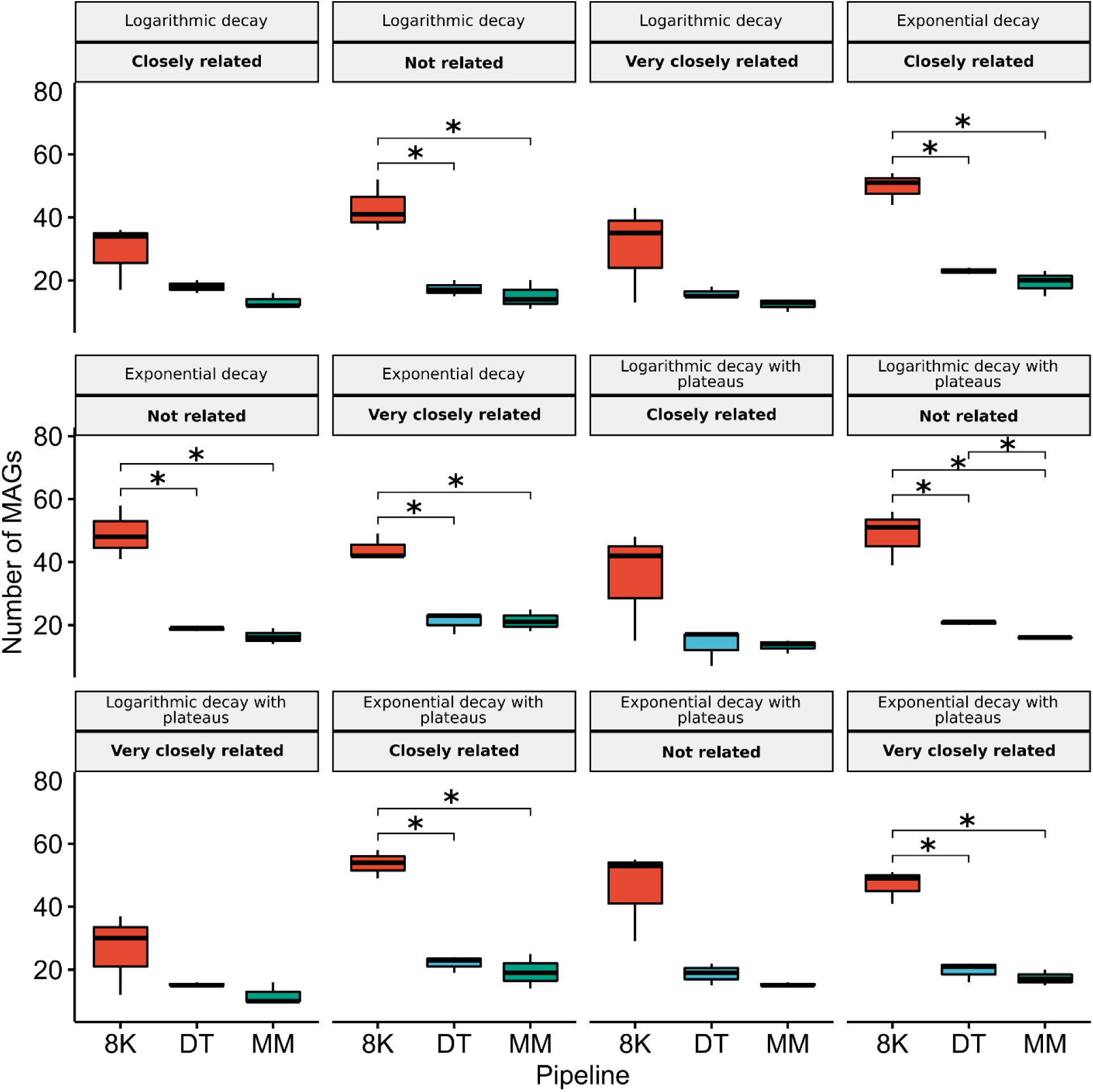
Student’s T-test comparing metagenome-assemble genome counts between all pipelines used (8K, DT and MM) according to Taxonomic distribution (Not related; Closely related; Very closely related) and evenness distribution (Logarithmic decay; Exponential decay; Logarithmic decay with abundance plateaus; Exponential decay with abundance plateaus). The sequencing depth of the communities was kept at 60 million reads. (*P-value < 0.05).

### 2.3 Sequencing depth as a major factor influencing MAG recovery

The 8K pipeline, on average, only recovered 25% of the 42-original species in each community profile (Supplemental_Table_S02.xlsx). Also, a significantly lower number of True Positives (TP) were found in “closely related” communities following a logarithmic decay when compared to those following an exponential decay with abundance plateaus (*t-*test, *p* < 0.05) (Supplemental_Table_S15.xlsx). No statistically significant differences were found in the number of TPs when comparing sequencing depths or taxonomic distributions in any of the evenness distribution profiles (Supplemental_Table_S16.xlsx and Supplemental_Table_S17.xlsx).

DT recovered, on average, the highest percentage of original species per community profile (45%) (*t-*test, *p* < 0.05) (Supplemental_Table_S02.xlsx). On average, the DT pipeline recovered 85% of the original species at least once (Supplemental_Table_S18.xlsx). In “closely related” communities, a significantly higher number of TPs was obtained when the communities followed an exponential decay evenness distribution compared to logarithmic decay (*t-*test, *p* < 0.05) (Supplemental_Table_S19.xlsx and Figure 3). A significantly lower number of TPs were observed in communities sequenced at 10 million reads than those sequenced at 60 million reads irrespective of evenness distribution (*t-*test, *p* < 0.05) (Supplemental_Table_S20.xlsx and Supplemental_Fig_S15.pdf).

**Figure 3.**
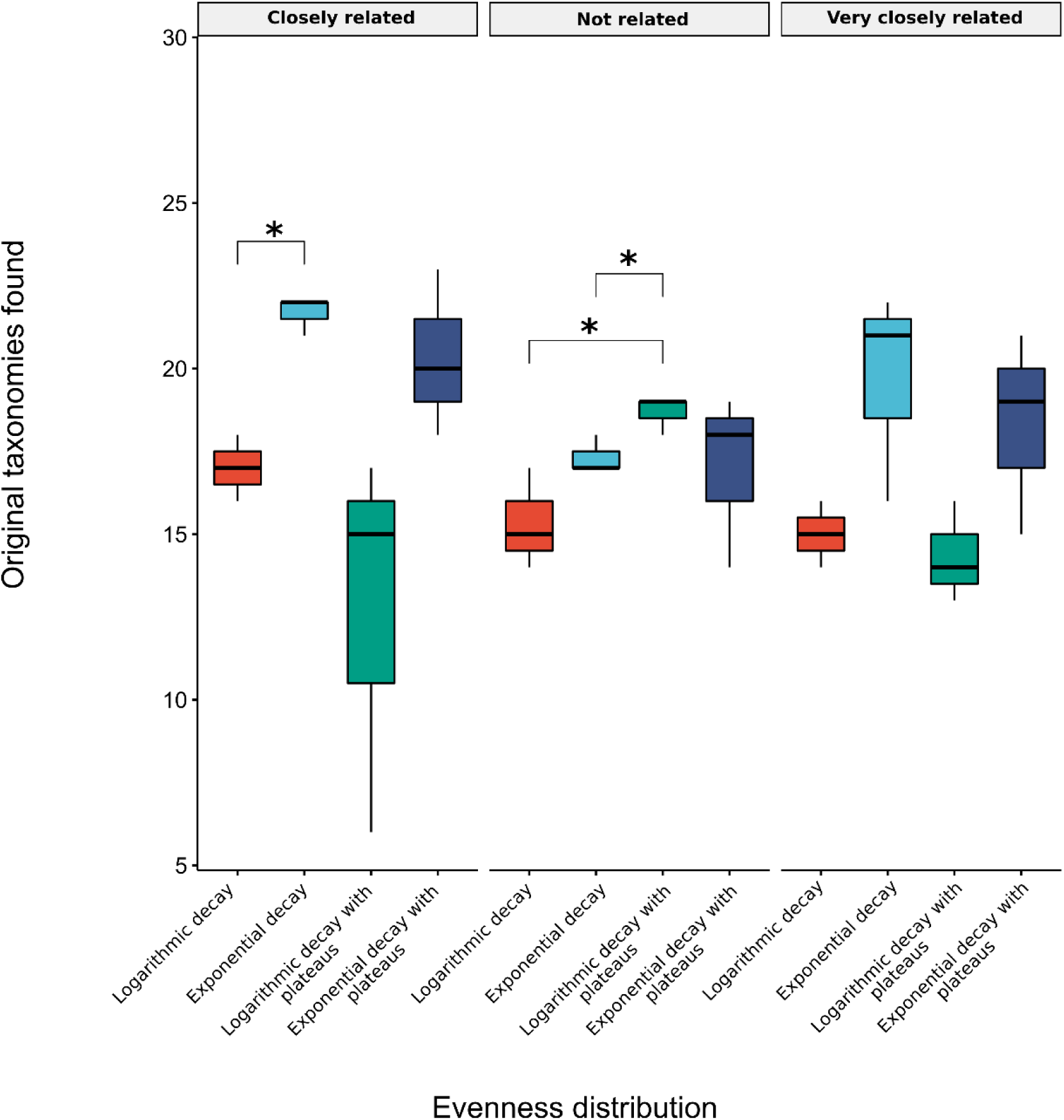
Student’s T-test comparing True Positives (Species recovered and in the original communities) obtained by the DT pipeline in communities following different evenness distributions (Logarithmic decay; Exponential decay; Logarithmic decay with abundance plateaus; Exponential decay with abundance plateaus) for each taxonomic distribution (Not related; Closely related; Very closely related). The sequencing depth of the communities was kept at 60 million reads. (*P-value < 0.05).

**Figure 4.**
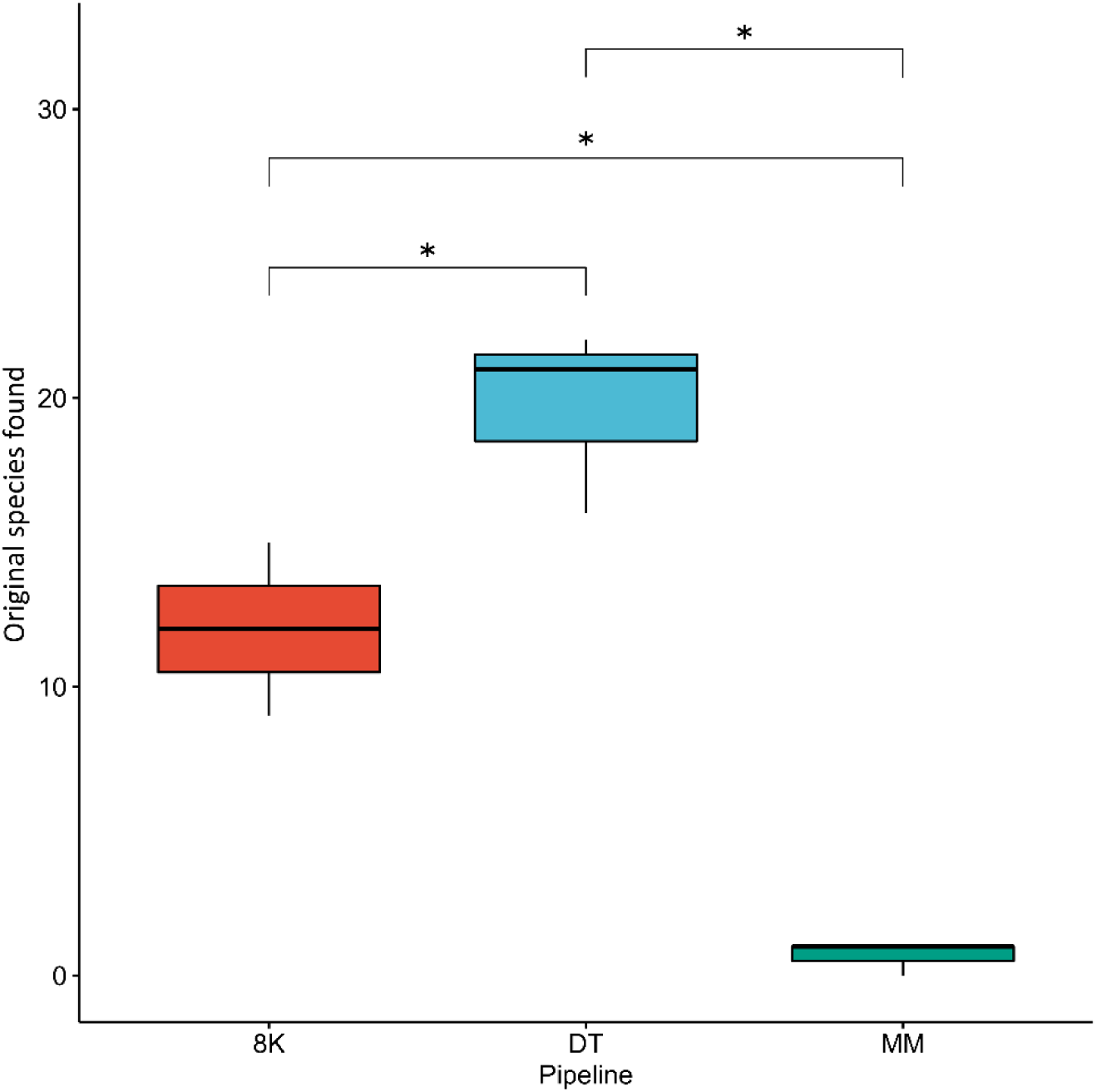
Student’s T-test comparing True Positives (Species recovered and in the original communities) etween all pipelines used (8K, DT and MM) in communities composed of species with equal abundance (i.e., equal evenness distribution). The sequencing depth of the communities was kept at 60 million reads. (*P-value < 0.05).

**Figure 5.**
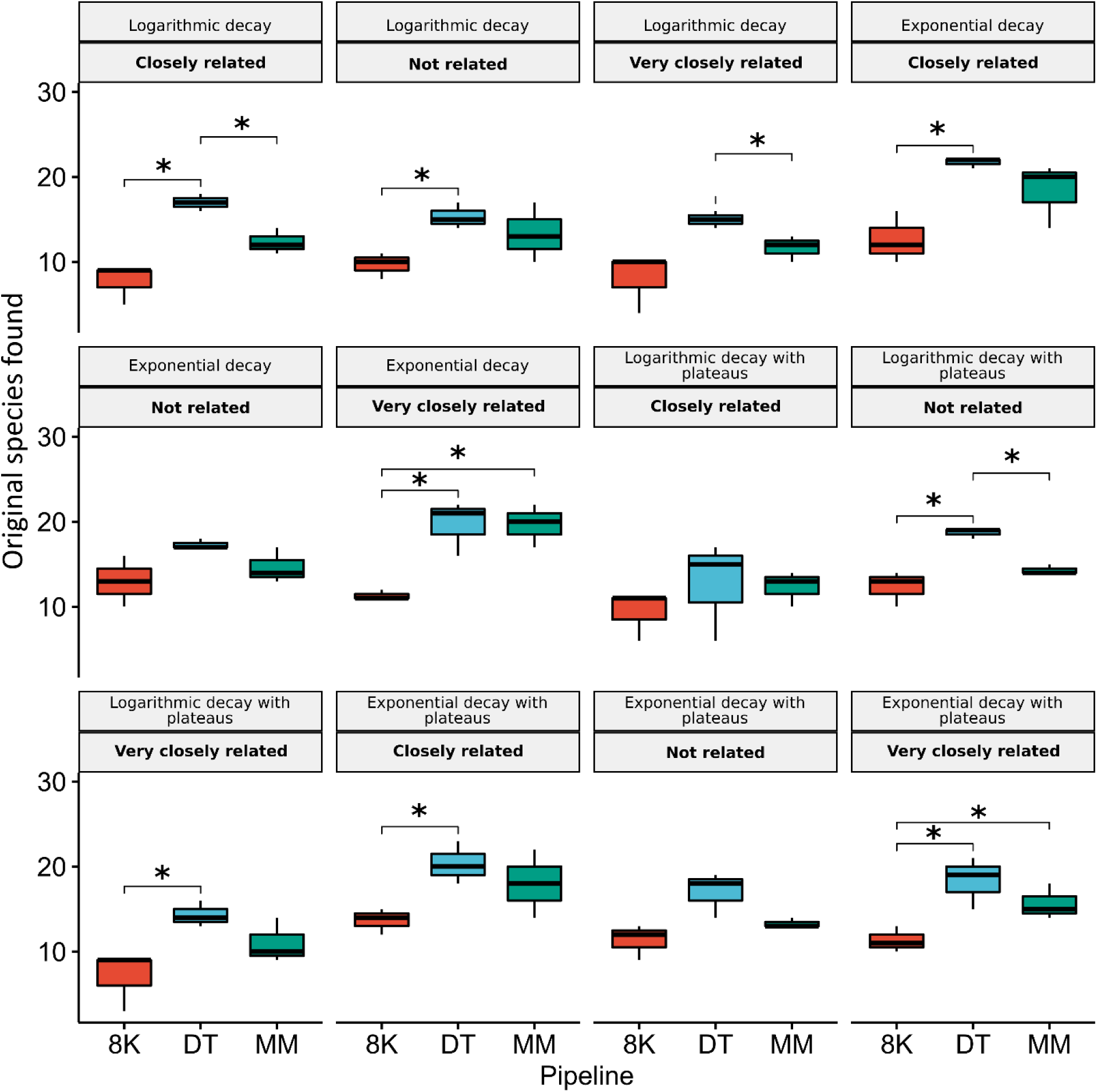
Student’s T-test comparing True Positives Species recovered and in the original communities) between all pipelines used (8K, DT and MM) according to Taxonomic distribution (Not related; Closely related; Very closely related) and evenness distribution (Logarithmic decay; Exponential decay; Logarithmic decay with abundance plateaus; Exponential decay with abundance plateaus). The sequencing depth of the communities was kept at 60 million reads. (*P-value < 0.05).

The number of TPs was significantly lower in communities following a logarithmic decay and sequenced at 10 and 30 million reads compared to 180 million reads (*t-*test, *p* < 0.05) (Supplemental_Table_S20.xlsx and Supplemental_Fig_S15.pdf). However, no significant differences were observed in the number of TPs recovered between communities sequenced at depths greater than 60 million reads.

The MM pipeline recovered, on average, 27.5% of the original species (Supplemental_Table_2.xlsx). In “very closely related” communities, a significantly higher number of TPs were observed under exponential decay evenness distributions when compared to those with logarithmic decay (*t-*test, *p* < 0.05) (Supplemental_Table_S21.xlsx and Supplemental_Fig_S16.pdf). A significantly higher number of TPs were observed in communities sequenced up to 30 and 60 million reads compared to 10 million reads (*t-*test, *p* < 0.05). However, communities sequenced at 120 and 180 million yielded a significantly lower number of TPs compared to communities sequenced below 60 million reads (*t-*test, *p* < 0.05) (Supplemental_Table_S22.xlsx and Supplemental_Fig_S17.pdf), irrespective of their evenness distribution. No statistically significant differences in the number of TPs were observed between communities of different taxonomic distributions in any of the evenness distributions (Supplemental_Table_S23.xlsx).

#### 2.3.1 DAS tool recovers the highest number of original species

The DT pipeline yielded a significantly higher number of original species than the MM and 8K pipelines, irrespective of sequencing depth, taxonomic relatedness and evenness distribution (*t-*test, *p* < 0.05) (Supplemental_Table_S24.xlsx – Supplemental_Table_S26.xlsx, Supplemental_Fig_S18.pdf and Figures 4-5).

The 8K pipeline also obtained a significantly lower number of TPs than the MM pipeline in communities following an exponential decay evenness distribution sequenced at 30 and 60 million reads (*t-*test, *p* < 0.05) (Supplemental_Table_S26.xlsx and Supplemental_Fig_S18.pdf). Inversely, the MM pipeline yielded a significantly lower number of TPs than 8K in communities following an exponential decay evenness distribution but sequenced at 120 and 180 million reads (*t-*test, *p* < 0.05) (Supplemental_Table_S26.xlsx and Supplemental_Fig_S18.pdf).

We did not identify contigs in the Prokaryotic MAGs originated from the Eukaryotic genomes added to the simulated microbial communities.

### 2.4 The highest number of false positives was found in communities with 60 million reads

The 8K, DT, and MM pipelines yielded an average of 8,8%, 8,1% and 9,4% of False Positives (FP) to the 42 species present in the original communities, respectively (Supplemental_Table_S02.xlsx). The 8K pipeline yielded a significantly higher number of FPs than the DT pipeline in 10 of the 23 community profiles (*t-*test, *p* < 0.05). The 8K pipeline also yielded significantly more FPs than the MM pipeline in 7 of the 23 community profiles (*t-*test, *p* < 0.05). These significant differences were observed in communities sequenced at 60 million reads. No clear pattern was observed when analyzing taxonomic relatedness and evenness distribution, suggesting that both parameters do not significantly influence the number of false positives.

#### 2.4.1 The MM pipeline yields the lowest number of False Positives but not unique species

The MM pipeline yielded the lowest of FPs (75) compared to the 8K (280) and DT (117) pipelines, irrespective of evenness distribution and sequencing depth (*t-*test, *p* < 0.05) (Supplemental_Table_S27.xlsx and Supplemental_Fig_S19.pdf). However, this difference was not reflected in the number of unique species. The number of unique species classified as FPs using the 8K, DT and MM pipelines was six, nine and eight, respectively. Furthermore, all three pipelines consistently obtained five FP species, *B. vaginale*, *B. pertussis*, *P. atlanticus*, *P. cladifontis* and *R. rhizogenes*. *D. amylolyticus* was recovered by both DT, and 8K pipelines whilst *B. burgdorferi*, *C. vibrioides* and *T. azollae* were recovered by DT and MM pipelines (Supplemental_Table_S04.xlsx).

### 2.5 DT pipeline displays the highest accuracy in species recovery

The DT pipeline displayed the highest average accuracy in species recovery (∼92% ±2.07) followed by the MM pipeline (∼88% ±15.57). The 8K pipeline displayed the least accuracy in species recovery (∼76% ±8.81) (Supplemental_Table_S02.xlsx). The DT pipeline was significantly more accurate than the 8K pipeline in 10 of the 23 community profiles, while the MM pipeline showed significantly higher accuracy rates than the 8K pipeline in 6 of the 23 community profiles (*t-*test, *p* < 0.05). The most significant differences in both cases (i.e., DT versus 8K and MM versus 8K) were observed in communities sequenced at 60 million reads. The results did not show a clear pattern in accuracy when analyzing evenness distribution and taxonomic relatedness.

### 2.6 Recovery of bacteria with linear and multiple chromosomes is not influenced by sample evenness, taxonomic relatedness or sequencing depth

As expected, none of the ‘special case’ genomes (i.e., linear or multi-chromosomes) belonging to fungi were recovered. All of the bacterial “special cases” were recovered at least once by each pipeline except *Borrelia burgdorferi* and *Brucella ovis*. This occured in communities where they were originally present in equal or higher abundances than other species (Supplemental_Table_S28.xlsx). The MM pipeline could not recover any of the ‘special case’ genomes in communities composed of species at equal abundances (Supplemental_Table_S04.xlsx). Interestingly, *B. melitensis* was recovered in the original communities and others irrespective of the pipeline used, the evenness and taxonomic distribution or sequencing depth (Supplemental_Table_S04.xlsx). On average, 62.5% (±13.5) of the bacterial ‘special cases’ were recovered across all community profiles (Supplemental_Table_S04.xlsx).

### 2.7 Detection of low abundant species is not influenced by sequencing depth

The average genome coverage and relative abundance of species across all communities, irrespective of evenness, taxonomy or sequencing depth, using the 8K pipeline was 103 (±137) and 0.0241 (±0.0258), respectively. From the original species recovered the archaea *Ferroglobus placidus* presented the lowest relative abundance (0.0003) and second lowest genome coverage (4) and was found in “random” community following a logarithmic decay evenness distribution and sequenced at 180 million reads. Interestingly, the archaea *Desulfurococcus amylolyticus* was wrongfully recovered in the same communities as in *F. placidus* with higher relative abundance (0.022) and coverage (549).

The average coverage and relative abundance of species across all communities, irrespective of evenness, taxonomy or sequencing depth. When we used the DT pipeline, the detection limit was 103 (±147) and 0.0267 (±0.0306), respectively, for genome coverage and relative abundance. From the original species recovered, the *Wolinella succinogenes* presented the lowest relative abundance (0.00046) and coverage (3) and was found in “very closely related” communities following a logarithmic decay evenness distribution and sequenced at 60 million reads. *Caulobacter vibriodes* was not present in the original set of species, but it was recovered in a community with the same profile with higher relative abundance (0.0821) and coverage (216). Interestingly, *Caulobacter vibriodes* also presented the highest coverage (1199) and relative abundance (0.1536) of the false positives when recovered in “random” communities following a logarithmic decay evenness distribution and sequenced at 180 million reads.

When we used the MM pipeline, the average genome coverage and relative abundance of species across all communities, irrespective of evenness, taxonomy or sequencing depth pipeline, was 88.3 (±98.4) and 0.03 (±0.0308), respectively. From the original taxa recovered, *Borrelia anserina* presented the lowest relative abundance (0.00073) and third lowest coverage (10x) and was found in” very closely related” communities following a logarithmic decay evenness distribution and sequenced at 60 million reads. *Granulicella mallensis* presented the lowest coverage (8x) from the original species included in the communities, but in “random” communities following an exponential decay evenness distribution and sequenced at 10 million reads. Of the taxa recovered and not included in the original species list, *C. vibriodes* presented coverage of 406x and the highest relative abundance (0.148). It was found in” closely related” communities following a logarithmic decay evenness distribution and sequenced at 60 million reads.

The relative abundances of all species (original and recovered) in each community profile using all pipelines are available in Supplemental_Table_S04.xlsx.

### 2.8 Close-related species have a low rate of recovery in all pipelines

In “closely related” communities, the 8K pipeline recovered on average 0.11% (±0) (Supplemental_Table_S29.xlsx) of the established “species pairs” (Supplemental_Table_S30.xlsx). In “very closely related” communities, only 0.076% (±0.04) of the established “species pairs” were recovered on average (Supplemental_Table_S28.xlsx). The 8K pipeline recovered the highest percentage of “species pairs” in “closely related” communities following an exponential decay with abundance plateaus evenness distribution (Supplemental_Table_S29.xlsx). The complete set of results of recovered “species pairs” is shown in Supplemental_Table_S31.xlsx.

In “closely related” communities, the DT pipeline recovered on average 0.31% (±0.09) (Supplemental_Table_S29.xlsx) of the established “species pairs” (Supplemental_Table_S30.xlsx). In “very closely related” communities, 0.26% (±0.01) of the established “species pairs” were recovered on average (Supplemental_Table_S28.xlsx). Following an exponential decay evenness distribution, the DT pipeline recovered the highest percentage of “species pairs” in “closely related” communities (Supplemental_Table_S29.xlsx). The complete set of results of recovered “species pairs” is shown in Supplemental_Table_S32.xlsx.

In “closely related” communities, the MM pipeline recovered on average 0.20% (±0.048) (Supplemental_Table_S29.xlsx) of the established “species pairs” (Supplemental_Table_S30.xlsx). In “very closely related” communities, only 0.26% (±0.041) of the established “species pairs” were recovered on average (Supplemental_Table_S29.xlsx). The MM pipeline recovered the highest percentage of “species pairs” in “closely related” communities following an exponential decay evenness distribution (Supplemental_Table_S29.xlsx). The complete set of results of recovered species-pairs is shown in Supplemental_Table_S33.xlsx.

The DT and MM pipelines recovered a significantly higher percentage of “species pairs” than the 8K pipeline in “very closely related” communities following a logarithmic decay evenness distribution (*t-*test, *p* < 0.05) (Supplemental_Table_S34.xlsx and Supplemental_Fig_S20.pdf).

## DISCUSSION

This study aimed to determine which factors influenced genome recovery from metagenomes, the detection limits of MAG recovery and which of the three binning tools performed best in MAG recovery. Additionally, we included fungi in our simulated microbial communities to determine their impact on the recovery of prokaryotic MAGs. Our prokaryotic MAGs showed no contamination from the genetic content of the Eukaryotic genomes indicating the presence of Eukaryotes does not impact the recovery of prokaryotic MAGs. Our results showed that MAG recovery was influenced by taxonomic relatedness and sequencing depth but not evenness distribution. For example, more MAGs were recovered by the DT pipeline in communities composed of closely related species than non-related species (following an exponential decay distribution) than by the 8K and MM pipelines. Sequencing depth did appear to influence MAG recovery in the DT and MM pipelines, but no clear pattern was observed. For example, the higher number of MAGs with increasing sequencing depths using the DT pipeline was not observed when using the MM pipeline, which depended on the combination of evenness distribution and sequencing depth. The MM pipeline uses differential binning coverage (Albertsen et al. 2013) (i.e., the abundance of species in the samples), which would lead to higher binning resolution with increased sequencing depths. However, our results aligned with studies evaluating sequencing depth (Rajan et al. 2019; Cattonaro et al. 2020), which suggested that higher sequencing depths do not necessarily translate to improved species recovery. Our results suggest a sequencing depth sweet spot of 60 million reads. Our tool comparison showed that the 8K pipeline always yielded more MAGs than the DT and MM pipelines, which may be linked to the use MetaBAT (Kang et al. 2015). MetaBAT combines tetra-nucleotide frequency and contig abundance probabilities during the binning process.

Another aspect of this study was to examine how much of the original species (True Positives) added to the simulated communities each pipeline could recover. Sequencing depth showed contrasting results between the DT and MM pipelines. Higher sequencing depths in the DT pipeline resulted in increased TPs, whilst the MM pipeline showed increased TP recovery in sequencing depths up to 60 million reads compared to 120 and 180 million reads. This result suggested that the combination of multiple binning tools used by the DT pipeline scales with sequencing depth contradicting the expected higher binning resolution with increased sequencing depths. Besides, using tools that rely on species abundances showed that higher sequencing depths did not result in the recovery of a higher number of unique taxa (Sieber et al. 2018). On this topic, Gweon and colleagues (Gweon et al. 2019) stated that relatively low sequencing depths were sufficient to capture the broad-scale taxonomic composition of samples.

All pipelines used in this study demonstrated, on average, a low ability to recover pairs of species from the same genus, indicating that MAG recovery is not the best technology to study closely related species in microbial communities. Our results corroborate those of Sevim and collaborators (Sevim et al. 2019), who suggested that genomes with higher sequence similarity lead to increased genome misassembly. However, their study did not consider the combination of species abundance, taxonomic relatedness and sequencing depth. Nevertheless, communities following an exponential decay with abundance plateau evenness distribution showed the highest ratio of species-pairs recovered irrespective of the pipeline. Further, the DT pipeline always yielded more TPs than the 8K and MM pipelines in any parameter combinations (evenness versus taxonomy, sequencing depth versus evenness distribution, equal abundance communities) which suggests that the combination of multiple binning tools is more accurate in characterizing community composition. Previous studies have suggested this increased accuracy due to the DT pipeline methodology (Sieber et al. 2018; Maguire et al. 2020).

In this study, we also aimed to determine how species not included in the simulated communities (False Positives) were recovered. None of our selected variables strongly influenced the number of FPs obtained by each pipeline, although the most significant differences were observed in communities sequenced at 60 million reads. However, the 8K pipeline always yielded more FPs than the DT and MM pipelines, independent of the community profile. Seven of the nine unique FP could be explained by misannotations since they belonged to the same genus as the original species. For example, *B. pertussis* is a Gram-negative bacterium that causes whooping cough (Vásconez Noguera et al. 2021) and belongs to the same genus as *B. bronchiseptica*, initially included in community A. In the case of *B. vaginale*, a Gram-positive bacterium that colonizes the vagina and contributes to the vaginal microbiome, studies have shown that bifidobacteria from vaginal and gut microbiomes are indistinguishable using comparative genomics (Freitas and Hill 2018). The two remaining FPs shared the same family as the three original species. For example, *P. caldifontis* is a thermophilic Gram-negative bacterium commonly found in hot springs (Mori et al. 2014) and shares the same family as *P. thermarum* (included in the original communities). Because of these results, once important species are identified in the MAGs recovered, we suggest additional molecular tools such as qPCR or fluorescence in situ hybridization (FISH) should be employed to validate MAG recovery results (Singleton et al. 2021; Ma et al. 2022).

We observed the influence of sequencing depth in communities following a logarithmic decay evenness distribution. Such communities (e.g., bioreactors) comprised species at higher abundances (i.e., dominant) than most of their counterparts. Thus, increasing sequencing depth will lower the detection limit allowing the recovery of low-abundant species. Inversely, communities following an exponential decay evenness distribution (e.g., pristine communities) may comprise a higher number of species at similar abundances and a lesser fraction at low abundances. Trying to lower the detection limit by increasing sequencing depth will have little influence on the recovery of more species. The increase in sequencing depth may also increase the coverage and relative abundance of recovered species. However, the fact that some FPs present coverages and relative abundance greater than TPs by several folds suggests that robust community identification should not exclusively depend on these measures.

Of the 11 special cases (i.e., bacteria with linear or multiple chromosomes) added to the mock communities, only *Borrelia burgdorferi* and *Brucella ovis* were not recovered. The non-recovery of *Borrelia burgdorferi* may be linked to a misclassification rather than its linear genome since we did recover *Borreliella burgdorferi* whose genome is also linear. The misclassification can also be tied to dividing the genus *Borrelia* into two genera which have sparked some debate (Gupta 2019). Similarly, the absence of *B. ovis* in our recovered MAGs may also be linked with misclassification due to high sequence similarities. Tsolis and colleagues (Tsolis et al. 2009) have shown that *B. ovis* shares similar chromosome sizes (bp), GC content and number of protein-coding genes to other Brucella species such as *B. melitensis* (recovered in our data). Our data are insufficient to assess the influence of linear or multi-chromosome genomes on MAG recovery but suggest it may increase the challenge of taxonomic classification. However, current methods for MAG recovery do not address this issue. Thus, it would be beneficial that future studies attempt to identify signatures that discriminate between linear and multiple chromosomes in MAGs.

While this study focused only on prokaryotes, similar studies of MAG recovery of Eukaryotes and viruses (DNA and RNA) may help improve ecological and biotechnological insights generated from genome-centric metagenomics.

Characterizing microbial communities is a challenging task due to their complexity. Our study evaluated the combined effect of species abundance, taxonomic proximity between species and sequencing depth in prokaryotic MAG recovery. Our analyses suggested that none of the variables used (species abundance, taxonomic proximity and sequencing depth) is solely responsible for MAG recovery or high accuracy in community characterization. However, sequencing depth and taxonomic proximity between species do appear to have a stronger influence on MAG recovery than sample evenness. The point was supported by the very low percentages of taxonomically related “species pairs” recovered independently of the community profile or pipeline used. Integrating multiple binning tools via a consensus approach appears to perform better. Furthermore, our data indicated that coverage and relative abundance might not be the main drivers of MAG recovery since species with high abundances in the original communities did not necessarily guarantee their recovery. Additionally, the recovery of false positives with relative abundances and coverages many times greater than those of the original species suggests that species abundance is unreliable in characterizing a microbial community accurately. Interestingly, the difference in coverages for all “species pairs” recovered followed the same trend as their initial abundances in almost all communities, suggesting a slight connection between species abundance and genome coverage. Nevertheless, further studies need to be carried out to confirm this hypothesis. Genome linearity or multi-chromosomes had no visible effect on MAG recovery in our study. However, to fully determine if this is the case, a more extensive study should be performed, including a larger number of genomes.

The pipeline used is also crucial since accuracy can range from 76% (when using the 8K pipeline) up to 92% (when using the DT pipeline). While the high accuracy obtained by the DT pipeline is promising, it is necessary to consider how the error rates become relevant for large MAG datasets. For every 10000 MAGs recovered, the number of false positive MAGs can range from 800 to 2400, depending on the pipeline. Therefore, our study demonstrated the need to verify findings based on MAGs, especially when novel species or special metabolic functions are highlighted.

## METHODS

### Workflow

The workflow for this study is shown in Figure 6, which comprised three main sections, pre-processing, binning and post-processing.

**Figure 6.**
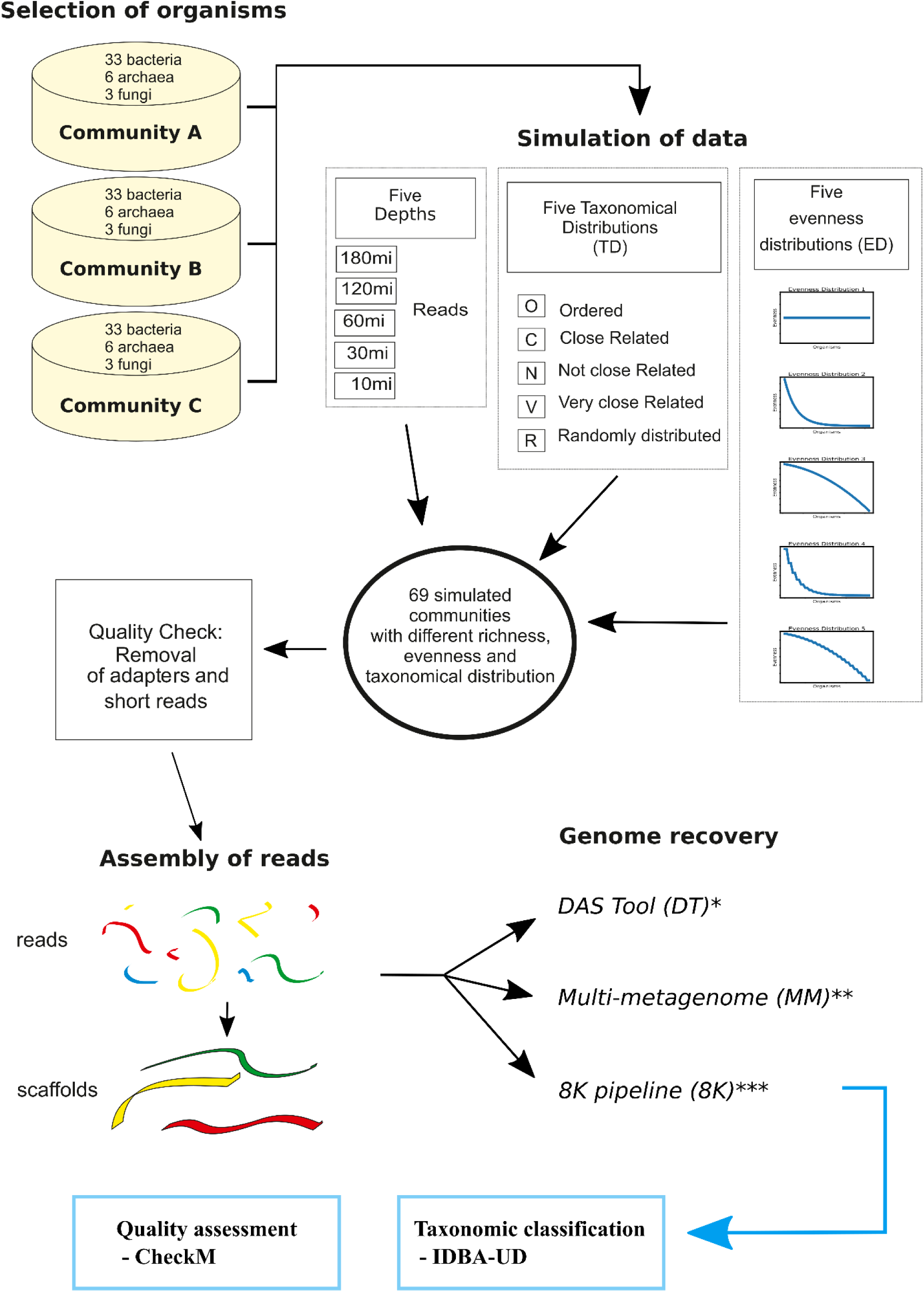
Workflow used in this study. First, we proceeded with species selection and sequence retrieval from the National Center for Biotechnology Information (NCBI). Next, community profiles were generated based on species abundance, taxonomic distribution and sequencing depth. Metagenomes were simulated for each community profile using MetaSim. A quality check was performed to remove adapters and short reads. The next step consisted in assembling reads into scaffolds and performing post-assembly quality checks. For genome recovery, three pipelines were used: DAS Tool (DT) (Sieber et al. 2018), Multi-metagenome (MM) (Albertsen et al. 2013) and the pipeline used to recover more than 8000 metagenome-assembled genomes (MAGs) (8K) (Parks et al. 2017). Completeness and contamination of MAFs was assessed using CheckM (Parks et al. 2015). Taxonomic classification of the MAGs was performed by IDBA-UD (Peng et al. 2012).

### Pre-processing

#### Selection of species of the in silico microbial communities

A total of 126 species were selected to represent the three main branches in the tree of life: 99 bacteria, 18 archaea and nine fungi (Supplemental_Table_S35.xlsx). Species were selected if a complete genome and the raw sequencing data were available in the National Center for Biotechnological Information (NCBI). Additionally, species were selected so that each community included taxonomic groups consisting of three species belonging to the same family and two of them to the same genus. We divided these 126 species into three groups of 42 (33 bacteria, six archaea and three fungi). A total of 12 taxonomic sets of species were present in each community. Next, each set of species was used to assemble communities with varying degrees of species abundance (evenness), taxonomic relatedness and sequencing depth.

#### Establishing taxonomic relatedness

The taxonomic relatedness of the species in this study were used to create profiles of taxonomic distribution. The species were grouped in triplets with different taxonomic distributions. First, a species was randomly set as the reference for a given phylum. Species belonging to the same genus as the reference were defined as “Very closely related” while species belonging to the same family (but not genus) as the reference were defined as “Closely related” (Supplemental_Table_S30.xlsx). For example, the first triplet *Acetobacter aceti* would be randomly selected from the phylum Proteobacteria. The genome considered “Very closely related” would be *Acetobacter persici* since it belongs to the same genus. The genome considered “Closely related”, would be *Gluconacetobacter diazotrophicus* since it belongs to the same family but not genus. The same procedure would be applied to all other genomes in each dataset. Communities were constructed according to five relationships between organisms: “Ordered”, “Random”, “Very closely related”, “Closely related” and “Not closely related” (Supplemental_Table_S36.xlsx). In the “Ordered” communities, species were simply placed based on their triplets. The “Random” communities were constructed without any specific criteria. The “Very closely related” communities were constructed considering their relationship with each other (e.g. species belonging to the same genus). Similarly, the “Closely related” communities were constructed considering the relationship between species at the family level. Lastly, “Not closely related” communities were constructed, so species with closely related relatives were not added at similar abundances. Examples of community definitions and relationships are shown in Supplemental_Table_S36.xlsx. These community definitions and relationships did not include fungi and special cases of bacteria genome topology (consisting of linear or multiple chromosomes). However, representatives of species with linear (*Vibrio cholerae* and *Rhodobacter sphaeroides*) and multiple chromosomes (*Bacillus subtilis*, *Streptomyces griseus*, *Borrelia burgdorferi*, *Paracoccus denitrificans*, *Planktomarina temperate*, *Vibrio fluvialis*, and *Brucella melitensis*) were included in each simulated community. The list of all species used per community and type of chromosome is available in Supplemental_Table_S35.xlsx.

#### Evenness and distribution profiles

The literature describes abundance and species phylogenies as the main parameters controlling MAG recovery (Papudeshi et al. 2017). In this study, we define community evenness as a function of abundance distribution and distribution arrays, i.e., which species occupy different levels of abundance in the community (Equation 1).

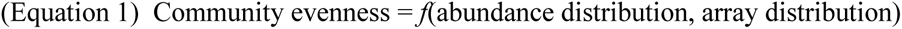

Evenness refers to the relative number of organisms of each species inside a given environment (system), which, in this case, is the main degree of freedom that will vary amongst the different communities. In a metagenomic community, evenness and abundance can sometimes be used interchangeably. The abundance of genomic data can be evenly distributed (equal evenness), normally or disruptively. In this study, we simulated abundance in several different states of evenness distribution (equal evenness, logarithmic decay, exponential decay, logarithmic decay with abundance plateaus and exponential decay with abundance plateaus. Communities classified as logarithmic decay with abundance plateaus are composed of species whose abundance follows a logarithmic decrease but with intervals of equal abundances for pairs of species. Similarly, communities classified as exponential decay with abundance plateaus are composed of species whose abundance follow an exponential decrease but with intervals of equal abundances for pairs of species (Supplemental_Fig_S21.pdf). Additionally, we coupled different abundance profiles with different taxonomic distribution profiles.

#### Sequencing depth

To evaluate the influence of sequencing on the recovery of MAGs, we simulated communities with five sequencing depths (10, 30, 60, 120 and 180 million paired-end ILLUMINA reads, 2×150bp).

#### In silico metagenomic library preparation

We generated 23 libraries for each group of species (hereafter, A, B and C): 12 through the combination of four abundance distribution profiles (logarithmic decay, exponential decay, logarithmic decay with abundance plateaus and exponential decay with abundance plateaus) with three taxonomic distribution profiles (“Very closely related “, “Closely related” and “Not closely related”) at a depth of 60 million reads. Ten libraries were generated combining the logarithmic decay and exponential decay profile with all the possible sequencing depths (10, 30, 60, 120 and 180 million reads) with no specific criteria for the species taxonomic relatedness (“Random” taxonomic distribution profile). A single library was generated with all species at equal abundances, “Ordered” taxonomy and a sequencing depth of 60 million reads. This library was used to test MAG recovery at identical species abundances.

### Processing

#### Simulation of High-throughput sequencing data

The reads were generated using the MetaSIM Sequencing simulator (v0.9.1) (Richter et al. 2008). The fragment size was set to 180 bp with a read size of 125 bp.

#### Assembly

The reads were assembled into scaffolds of each simulated metagenomic library using IDBA-UD (Peng et al. 2012) with default parameters.

#### Draft genome reconstruction pipeline

We used three pipelines to recover high-quality metagenome-assembled genomes (MAGs). The multi-metagenome (MM) pipeline (Albertsen et al. 2013) assembles near-complete draft genomes in a two-step process (i.e., primary binning is performed independently of sequence composition followed by refinement of population genomes using sequence composition-dependent methods and visualization tools). The DAS Tool (DT) pipeline (Sieber et al. 2018) generates bins by integrating multiple binning algorithms. The pipeline developed by Parks and co-workers (8K) (Parks et al. 2017) uses MetaBAT (Kang et al. 2015) and integrates genome abundances and tetranucleotide frequencies to generate bins.

#### Bin quality assessment and filtering

Completeness and contamination measures were obtained via CheckM (Parks et al. 2015). Bin quality was determined using the approach by Parks and colleagues (Parks et al. 2017) (Equation 2). Only bins with a quality score above 50 were considered for subsequent analyses.

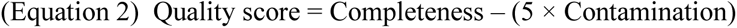

#### Assessing the performance of binning pipelines

We classified each genome used in the different communities with GTDB-tk version 1.3 (Chaumeil et al. 2020) to determine the performance of each pipeline and the influence of species abundances, taxonomic relatedness and sequencing depth. True positives (TP) were defined as MAGs correctly classified in each community, false positives (FP) were defined as MAGs with taxonomy not included in the original communities, and false negatives (FN) were defined as MAGs present in the original community but not found after the recovery process. Additionally, to test the effect of closely and very closely related species at equal or similar abundances on genome recovery, we calculated the fraction of the pairs of species that were recovered in the same community. We determined the accuracy of MAG recovery using Equation 3:

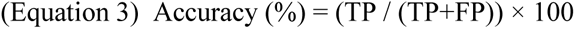

#### Detection limits

Usually, the number of species recovered in a sample using metagenomics is a product of the number of reads sequenced (Ebinger et al. 2021). The higher the number of reads, the more likely we can recover species at low quantities. We calculated each MAG’s coverage (Equation 4) and relative abundance (Equation 5) in their respective library to test this hypothesis.

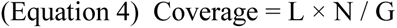

In equation 4, L is the mean length of the reads per library, N is the total number of mapped reads in the MAG, and G is the size of the MAG (in base pairs).

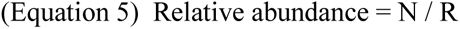

In equation 5, N is the total number of mapped reads in the MAG, and R is the total number of reads in a sample.

We also tested whether there is a correlation between relative abundance and coverage and the number of false positive species (i.e., species recovered but not in the original communities).

#### Statistical analysis

Student’s t-test was used to determine significant differences obtained in the number of MAGs, TPs and FPs when comparing sequencing depth, taxonomic relatedness and evenness distribution profiles. Results were considered statistically significant when the P values were below 0.05.

#### Computational resources

All *in silico* operations were performed using a High-Performance Computer (HPC). The HPC cluster was composed of 44 nodes with 28 cores each. Each node had 224 GB of main memory usable for jobs.

## DATA ACCESS

The true positive metagenome-assembled genomes (MAGs) (4638 out of 5217) obtained in this study are available at the National Center for Biotechnology Information (NCBI) repository with the sample accession numbers SAMN34004744 - SAMN34012094 (BioProject PRJNA950613). The complete set of MAGs (including the false positives) and simulated metagenomic libraries are available on the long-term data archive at the Helmholtz Center for Environmental Research – UFZ data center using the link (https://www.ufz.de/record/dmp/archive/13476).

## ACKNOWLEDGMENTS

This work was funded by the Helmholtz Young Investigator grant VH-NG-1248 Micro’ Big Data’ and the Deutsche Forschungsgemeinschaft (DFG, German Research Foundation) – project number 460129525. JK was supported by São Paulo Research Foundation (FAPESP; grant 2019/03396-9 and 2022/03534-5).

## AUTHOR CONTRIBUTIONS

UNR conceptualized, coordinated, and supervised all work. JPS contributed to the conceptualization of and supervision of all work. RBT downloaded all original genomes, generated all simulated microbial communities, and performed Metagenome-Assembled Genome (MAG) recovery and taxonomic classification of all MAGs. JCK analyzed the recovery of genome pairs and relative abundances and coverages of all MAGs. JPS performed all genome recovery and statistical analysis. SM and DS provided an essential critical review of the manuscript. All authors read and approved the final manuscript.

